# A molecular switch at the yeast mitoribosomal tunnel exit controls cytochrome *b* synthesis

**DOI:** 10.1101/2025.01.30.635641

**Authors:** Andreas Carlström, Joseph B. Bridgers, Mary Couvillion, Abeer Singh, Ignasi Forné, Axel Imhof, L. Stirling Churchman, Martin Ott

**Author notes:** These authors contributed equally to this work (Andreas Carlström, Joseph B. Bridgers). Correspondence (S.C.), (M.O.).

## Abstract

Mitochondrial gene expression needs to be balanced with cytosolic translation to produce oxidative phosphorylation complexes. In yeast, translational feedback loops involving lowly expressed proteins called translational activators help to achieve this balance. Synthesis of cytochrome *b* (Cyt*b* or *COB*), a core subunit of complex III in the respiratory chain, is controlled by three translational activators and the assembly factor Cbp3-Cbp6. However, the molecular interface between the *COB* translational feedback loop and complex III assembly is yet unknown. Here, using protein-proximity mapping combined with selective mitoribosome profiling, we reveal the components and dynamics of the molecular switch controlling *COB* translation. Specifically, we demonstrate that Mrx4, a previously uncharacterized ligand of the mitoribosomal polypeptide tunnel exit, interacts with either the assembly factor Cbp3-Cbp6 or with the translational activator Cbs2. These reciprocal interactions determine whether the translational activator complex with bound *COB* mRNA can interact with the mRNA channel exit on the small ribosomal subunit for translation initiation. Organization of the feedback loop at the tunnel exit therefore orchestrates mitochondrial translation with respiratory chain biogenesis.

## Introduction

Organellar protein synthesis is a remnant of the endosymbiotic origin of mitochondria and chloroplasts. Proteins encoded in mitochondrial DNA (mtDNA) make up a small, but critical fraction of large multi-subunit assemblies, namely mitochondrial ribosomes (mitoribosomes) and the complexes driving oxidative phosphorylation (OXPHOS). Most subunits of the mitoribosome and OXPHOS are encoded in the nuclear genome, synthesized in the cytosol and imported into mitochondria, where they are assembled with the mitochondrial translation products. The dual genetic origin of OXPHOS subunits and their assembly into complexes with defined stoichiometry necessitates the coordination of both genetic systems for a balanced translational output ^1,2^. Expression of nuclear-encoded genes is tightly regulated by transcription factors, but organellar gene expression also can be modulated ^3^. While chloroplast gene expression employs a rather wide variety of options ^4–6^, accumulation of mitochondrial translation products is mainly adjusted through translational control ^7–9^ or through degradation of proteins produced in excess ^10–12^.

In the baker’s yeast, *Saccharomyces cerevisiae*, translational control of a subset of mitochondrial mRNAs is achieved through feedback regulation ^7,9,13^. By this, translational output is adjusted to levels that allow efficient assembly of the newly synthesized proteins. Here, the case of mitochondrial encoded cytochrome *b* (Cyt*b*) is best understood. Cyt*b* is the central subunit of the respiratory chain complex III and a highly hydrophobic membrane protein that participates in electron transport via its two heme *b* cofactors. Cyt*b* is assembled with nine nuclear encoded proteins to form complex III ^14^, which in turn associates with complex IV to form respiratory supercomplexes ^15,16^.

Synthesis of Cyt*b* by mitoribosomes depends on a set of translational activators (TAs), proteins that bind to the Cyt*b*-encoding mRNA (*COB*) to support translation. How exactly the TAs of *COB* mRNA (Cbs1, Cbs2 and Cbp1) work to activate translation of their client mRNA is currently not well understood. Proximity mapping has shown that *COB* TAs are dual localized on the surface of the mitoribosome, binding close to the mRNA channel exit (MCE) on the small ribosomal subunit (SSU) to aid translation initiation of the mRNA, and relocating to the polypeptide tunnel exit (PTE) of the large ribosomal subunit (LSU) upon translational repression ^17,18^. The localization of *COB* TAs to alternative sub-ribosomal locations is coincident with localization dynamics of the assembly factor Cbp3-Cbp6. This heterodimer localizes to the PTE ^18,19^ where it is able to interact with the newly synthesized Cyt*b*. Binding to Cyt*b* releases Cbp3-Cbp6 from the mitoribosome ^19^, after which the Cyt*b*-Cbp3-Cbp6 complex is channeled into the assembly line of complex III ^20,21^.

Localization of Cbp3-Cbp6 to the PTE is the key detection step of the *COB* mRNA feedback loop, which subsequently leads to translation activation. Its PTE association reports that the complex is available to chaperone and assemble newly synthesized Cyt*b*. In the event of stalled assembly, for example due to a deficiency in nuclear-encoded subunits, Cbp3-Cbp6 is sequestered in assembly intermediates and cannot activate translation. To this end, Cbp3-Cbp6 and the TA complex formed by Cbs1, Cbs2 and the *COB* mRNA localize to the PTE in a mutually exclusive manner ^17^, but how these reciprocal interactions are orchestrated and how these presumable dynamic interactions are conveyed to translational regulation remained unclear.

Here, we reveal the molecular mechanism underlying this translational feedback loop, which hinges on the dynamic, context-dependent interactions of TAs and Cbp3-Cbp6 at strategic sites on the mitoribosome. At the center of this regulatory circuit is Mrx4, a novel ligand of the mitoribosomal PTE that acts as a molecular switch. Mrx4 orchestrates the feedback loop by providing a mutually exclusive binding site for either the *COB* TA Cbs2 or the assembly factor Cbp3-Cbp6. This reciprocal binding to Mrx4 toggles between repression and activation of *COB* mRNA translation. Mrx4 represses translation of *COB* through sequestration of the TA complex at the PTE. Following the binding of Cbp3-Cbp6 to Mrx4, the TA complex is released from the PTE and relocates to the MCE for translation initiation. Thus, Mrx4 directly couples complex III assembly status to *COB* translation, ensuring precise coordination between mitochondrial and nuclear gene expression.

## Results

### Cbs1 and Cbs2 bind to the mitoribosome during early stages of translation

The polypeptide tunnel exit (PTE) and the mRNA channel exit (MCE) on the mitoribosome are two sites particularly important for organizing protein biogenesis and translational regulation, respectively (**Fig 1a**). Protein biogenesis factors like membrane insertases and chaperone-like assembly factors bind to the PTE to guide the nascent chain into the membrane and support protein folding ^22–25^, as also observed in other translation systems ^26,27^. Translational activators (TAs), each specific for one mRNA, bind to the MCE found on the SSU, where a canyon-like structure, formed by proteins bS1m (Mrp51) and mS43 (Mrp1), serves as their binding platform ^18,28^. Regulation of *COB* mRNA translation entails three TAs, Cbp1, Cbs1 and Cbs2, which were genetically mapped ^29^ to bind specific portions in the 5’-untranslated region (5’-UTR) of the *COB* mRNA (**Fig 1b**). Translational control of *COB* mRNA involves reciprocal localization of either the chaperone Cbp3-Cbp6 or the TAs Cbs1 and Cbs2 in complex with *COB* mRNA to the PTE ^17^. We employed selective mitoribosome profiling (sel-mitoRP) ^28^ to identify the timing of engagement of Cbs1 and Cbs2 with mitochondrial ribosomes (**Fig 1c**). Like in the case of Cbp1 ^28^, sel-mitoRP showed that ribosomes complexed with Cbs1 and Cbs2 can also be engaged in translation of other mRNAs (**Fig 1d**), likely reflecting a state of translational repression of *COB* mRNA due to an activated feedback loop ^17^. Importantly, and in line with a specific function during translation initiation, we found that Cbs1- and Cbs2-bound mitoribosomes are engaged in translation of the 5’ end of the *COB* open reading frame (ORF), but not at later stages (**Fig 1e**). This demonstrates that Cbs1 and Cbs2 play important roles in initiation of translation but leave the ribosome during elongation. Likewise, protected fragments of the *COB* mRNA 5’-UTR were observed in Cbs1 sel-mitoRP around GC-rich regions located around –100, -340, -600 and –950 (**Fig 1f**). Similar signals were observed for 5’-UTRs of other mitochondrial encoded mRNA ^28^, suggesting that they represent the TA binding sites on the mRNA.

**Figure 1:**
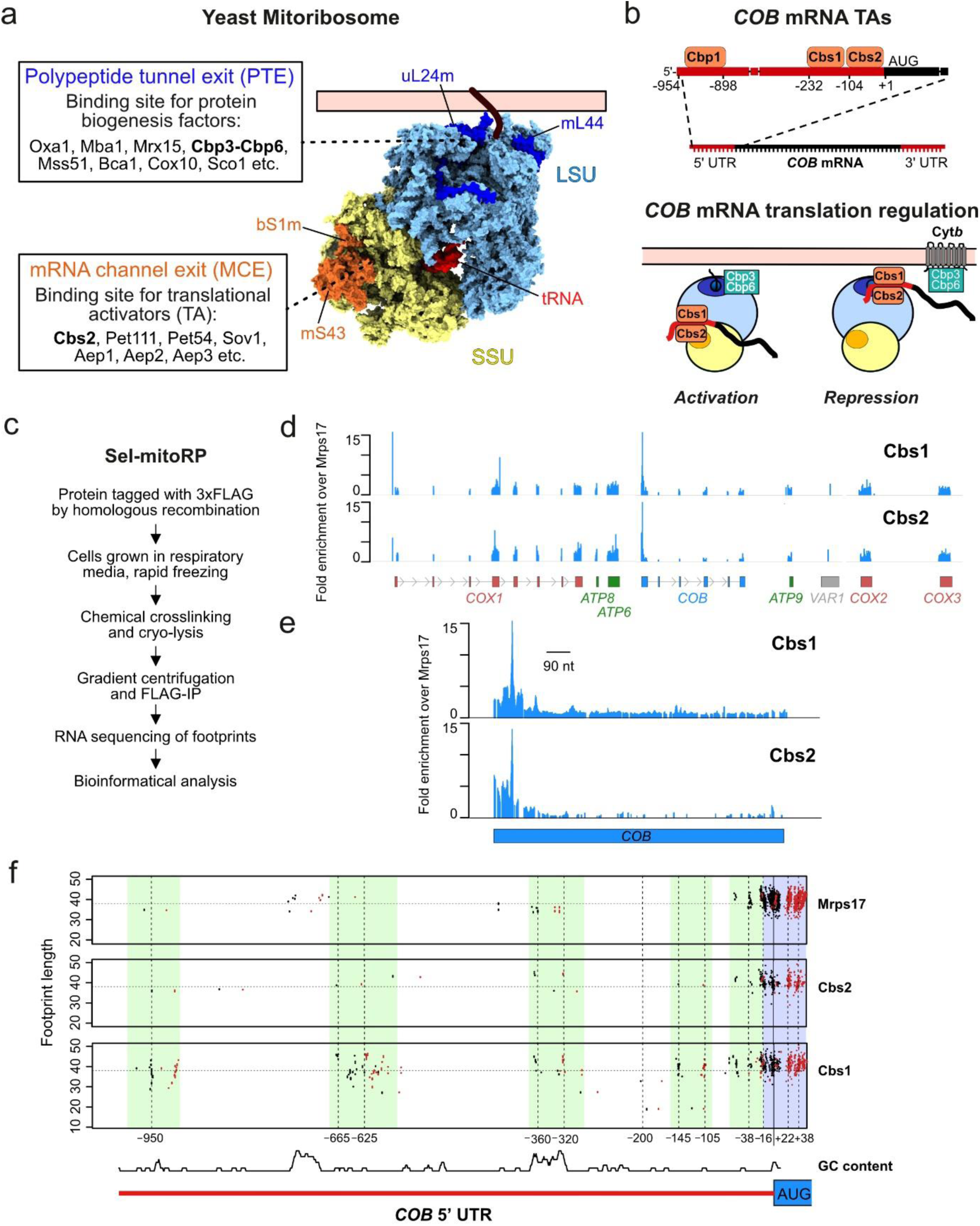
*COB* translational activators bind the 5’ UTR of *COB* mRNA and to the mRNA channel exit on the mitoribosomal SSU. **(A)** Illustration of the yeast mitochondrial ribosome (mitoribosome) highlighting two areas of importance for mitochondrial protein biogenesis. The polypeptide tunnel exit (PTE) on the large mitoribosomal subunit (LSU), highlighted with the two proteins uL24m and mL44, is the binding site for several factors involved in protein biogenesis and membrane insertion. The mRNA channel exit (MCE) on the small mitoribosomal subunit (SSU), highlighted with the two proteins bS1m and mS43, is the binding site for several proteins involved in activation of translation (TAs). Mitoribosome PDB ID: 5MRC ^30^. **(B)** Top: Schematic illustrating the estimated binding regions for the *COB* translational activator proteins Cbp1, Cbs1 and Cbs2 on the 5’-UTR of the *COB* mRNA. Bottom: The *COB* TAs are proposed to shuttle the *COB* mRNA between a translationally repressed state, bound close to the PTE, and active state where they bind at the mRNA channel exit for translation initiation. **(C)** General workflow for performing selective mitoribosome profiling (sel-mitoRP) of mitoribosomes crosslinked to and affinity-purified via specific protein factors. **(D)** Genome-wide view of protected mitochondrial RNA footprints after sel-mitoRP of mitoribosomes bound to Cbs1 and Cbs2. Enrichment of TA-specific reads relative to reads from mitoribosomes purified via Mrps17-3xFLAG are presented. **(E)** Sel-mitoRP of Cbs1- and Cbs2-bound mitoribosomes demonstrate an enrichment of footprints only in the 5’ portion of *COB* mRNA and during the start of translation. **(F)** V-plots of protected footprints at specified positions in the 5’-UTR of *COB* mRNA after sel-mitoRP of Cbs1- and Cbs2-bound mitoribosomes compared to non-selective mitoribosomes purified via Mrps17. Regions with marked footprints from the specific TAs are highlighted in green and footprints protected by the mitoribosome is highlighted in blue (-16 to +22). GC content: average GC content in 6 nt window (A/T=0, G/C=1), scale (0-1) not shown.

### Mrx4 is a novel PTE proximity interactor to Cbs2 and Cbp3

To further characterize the local environment of Cbs1 and Cbs2 in mitochondria, we utilized Bio-ID analyses with genomically integrated BirA*-tagged protein variants (**Fig 2a**) ^18^. By this, proteins in proximity of the bait proteins are labelled with biotin, which serves as a handle to purify these proximal proteins and determine their identity by mass-spectrometry. Cbs1 was found in proximity of the PTE protein mL44 but also the SSU protein uS12m (**Fig 2b**). Proximity mapping of Cbs2 revealed interactions with the LSU protein mL44, Sov1, a TA of *VAR1* mRNA^31^ which binds to the MCE of the SSU ^18^, and the uncharacterized protein Mrx4 (**Fig 2c**). Furthermore, in a Bio-ID experiment for Cbp3, Mrx4 was one of the most enriched proteins together with Cbp6, the mitoribosome receptor Mdm38 ^32^ and the complex III assembly factor Bca1/Fmp25 (**Fig 2d**). Mrx4 has previously been identified in proteomic analyses of mitochondrial ribosomes ^33^ and determined to be a high-confidence proximity interactor of the PTE ^18^, with a similar pattern as Cbp3 (**Fig 2e**).

**Figure 2:**
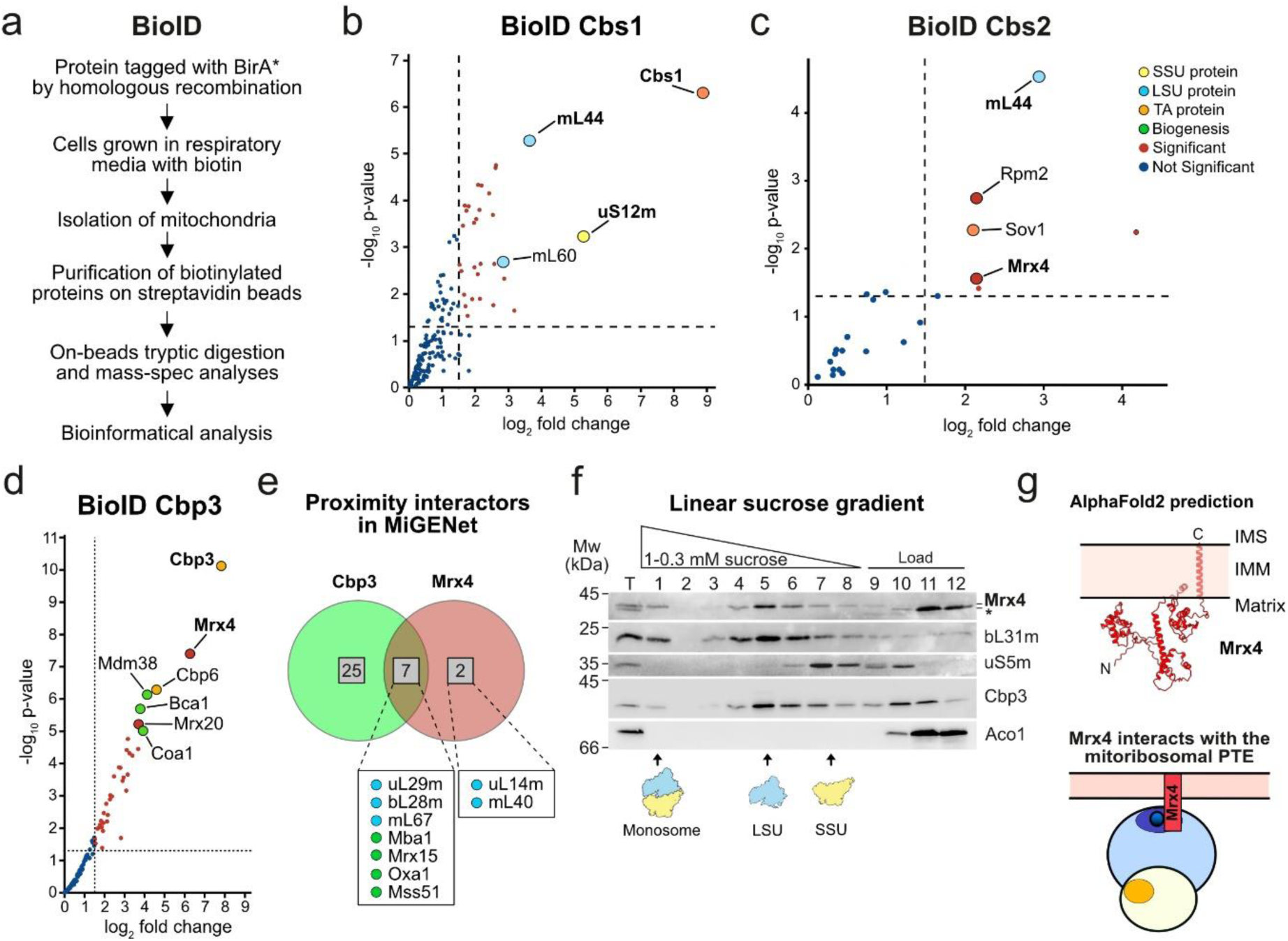
Mrx4 interacts with the PTE, Cbp3-Cbp6 and Cbs2. **(A)** Workflow of proximity labelling (Bio-ID) of mitochondrial proteins using a BirA*-tag genomically introduced to the C-terminus of target proteins. **(B)** Bio-ID of Cbs1 shows proximity to proteins of both the large (LSU) and small (SSU) mitoribosomal subunits. Data presented as log2 fold change compared to a Kgd4-BirA* control with a threshold set to 1.5. **(C)** Bio-ID of Cbs2 shows proximity to proteins of the LSU and SSU but also to the *VAR1* translational activator Sov1 and Mrx4. Data presented as log2 fold change compared to a Kgd4-BirA* control with a threshold set to 1.5. **(D)** Bio-ID of Cbp3 compared to the control Kgd4-BirA*, with a log2 fold change over 1.5 and a log10 p value under 0.5 considered significant. **(E)** First-neighbour analysis of common interactors of Cbp3 and Mrx4 extracted from the proximity interactome network MiGENet ^18^. **(F)** Linear sucrose gradient of a mitochondrial lysate shows that Mrx4 interacts quantitatively with the LSU (bL31m) but not with the SSU protein (uS5m) or matrix control Aco1. Cbp3 partially comigrates with the mitoribosome. (**G)** Top: The predicted structure of Mrx4 using *AlphaFold2* contains a single transmembrane (TM) helix at the very C-terminal end, with the majority of the protein located in the matrix with a loose fold consisting of three distinct domains. Bottom: Schematic showing Mrx4 interacting with Cbp3-Cbp6 at mitoribosomal PTE.

Likewise, Mrx4 co-migrated with the LSU on a linear sucrose gradient, an interaction that was not changed in strains lacking Cbp3 or Cbp1 (which leads to absence of *COB* mRNA) (**Fig 2f and S1a-b**). Similarly, deletion of Mrx4 did not affect the interaction of Cbp3 with the mitoribosome as evidenced by its co-migration with the LSU and the formation of a specific crosslinking product to the PTE protein uL29m (Mrpl4) (**Fig S1c-e)**. We found Mrx4 to localize to the mitochondrial matrix and to be anchored to the inner membrane with a single transmembrane helix (TM) at the C-terminus (**Fig 2g and S1f-i**). Interestingly, structure prediction using Alphafold2 suggested that Mrx4 is fairly unstructured with long flexible linkers connecting three more folded domains, which could be important for establishing protein-protein contacts (**Fig 2g**).

### Mrx4 mediates the feedback loop necessary for repression of *COB* translation

What is the functional relevance of Mrx4 interacting with the PTE? Given that Mrx4 is located in proximity of both Cbs2 and Cbp3, we hypothesized that it might be involved in the regulation of *COB* mRNA translation. Indeed, labelling of mitochondrial translation products with ^35^S-methionine *in vivo* demonstrated that absence of Mrx4 stimulated Cyt*b* synthesis, while absence of Cbp3 inhibited it (**Fig 3a**). Strikingly, in a double mutant lacking both Cbp3 and Mrx4, synthesis rates of Cyt*b* were restored (**Fig 3a**). To test directly whether Mrx4 is implicated in translational regulation, we utilized a reporter strain, where the *COB* ORF in mtDNA has been replaced with the coding sequence of *ARG8* (**Fig 3b**). Accumulation of Arg8, which is produced from an mRNA containing the 5′-UTR of *COB* mRNA, indicates how efficiently this mRNA can be translated, irrespective of whether the produced protein can be assembled into a functional respiratory chain ^19^. Deletion of *MRX4* did not decrease levels of Arg8, in contrast to deletion of *CBP3*, which abolished *cob::ARG8* translation (**Fig 3c**). Importantly, the decreased accumulation of Arg8 in the *cbp3Δ* strain was completely restored in the double mutant *cbp3Δmrx4Δ.* This effect was specific for Cbp3, as deletion of *MRX4* in cells lacking the other *COB* translational activators Cbs1 and Cbs2 could not restore Arg8 levels (**Fig 3c**). Likewise, deletion of the early complex III assembly factor Bca1 did not affect *cob::ARG8* translation and, subsequently, was not changed upon the deletion of *MRX4*.

**Figure 3:**
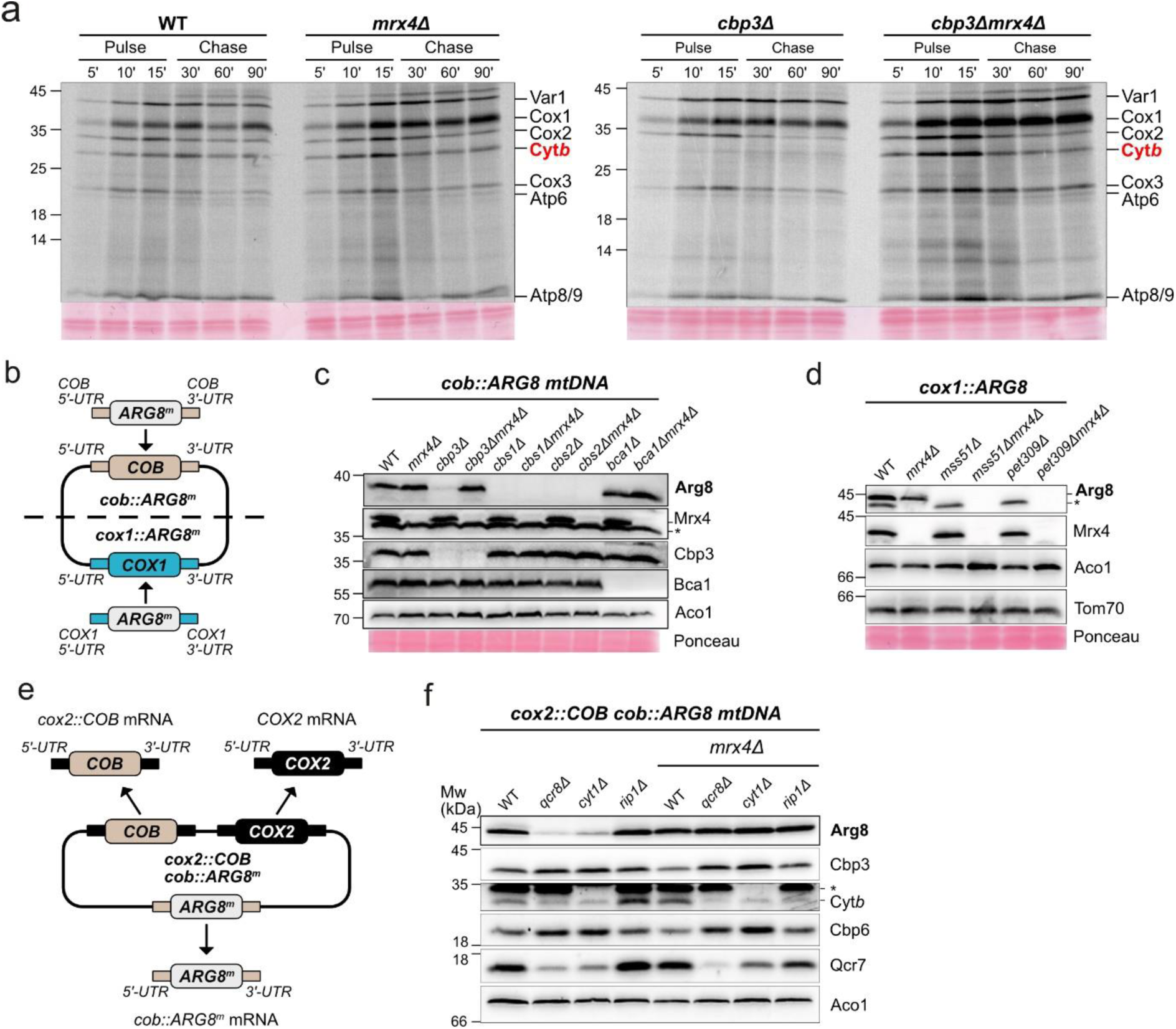
Mrx4 is needed for repression of the *COB* translational feedback loop. **(A)** *In-vivo* radiolabelling of indicated strains using ^35^S-methionine. Mitochondrial translation was followed for 15 min (pulse) and the stability of the synthesized proteins were monitored for 90 min (chase). Deletion of *CBP3* leads to a substantial decrease in Cyt*b* synthesis which is then restored again upon deletion of *MRX4*. **(B)** Schematic showing the mtDNA in the *cob::ARG8* and *cox1::ARG8* reporter strains. *ARG8* has replaced the open reading frames of *COX1* and *COB*, with the native 5’ and 3’ UTRs still intact. **(C-D)** Steady state levels of Arg8 and other indicated proteins in *cob:ARG8* and *cox1:ARG8* strains with deletions of *COB* and *COX1* translational activators together with deletion of *MRX4*. **(E)** Schematic showing the modifications of mtDNA in the *cox2::COB cob::ARG8* reporter strain. *ARG8* has replaced the open reading frame of *COB*, with the native 5’ and 3’ UTRs being present, while the native *COB* ORF has been engineered to be expressed under the control of *COX2* 5’ and 3’ UTRs. **(F)** Steady-state levels of Arg8 and other indicated proteins in strains containing the *cox2::COB cob::ARG8* mtDNA together with deletions of *QCR8*, *CYT1* and *RIP1* in combination with deletion of *MRX4*.

Next, we asked whether *MRX4* deletion could also reactivate translation of *COX1* mRNA, another example of a mitochondrially encoded mRNA that is regulated through a feedback loop ^34,35^. Strains lacking the *COX1* translational activators Mss51 or Pet309 in the analogous *cox1::ARG8* reporter system (**Fig 3b**) showed completely abolished accumulation of Arg8 (**Fig 3d**). Importantly, additional deletion of *MRX4* in the *mss51Δ* or *pet309Δ* strains did not restore Arg8 levels, revealing that Mrx4 plays a dedicated, direct role for Cbp3-Cbp6-dependent translation of *COB* mRNA.

Cbp3-Cbp6 shuttles between binding to newly synthesized Cyt*b* or to the mitoribosome, which leads to a new round of *COB* translation ^20^. Absence of nuclear-encoded, imported complex III subunits, like Qcr7, Qcr8 or cytochrome *c*_1_ (Cyt1), stalls complex III assembly and sequesters Cbp3-Cbp6 in early assembly intermediates. This translational repression can be clearly demonstrated in a strain where the native *COB* ORF is replaced by *ARG8*, while a second engineered *COB* ORF is expressed from an mRNA flanked by the *COX2* UTRs (**Fig 3e)**^20^. This results in the constitutive expression of *COB* by the *COX2* TAs, while *ARG8* serves as a reporter for translational efficiency of the *cob::ARG8* mRNA. Analysis with Western blot (**Fig 3f**) showed markedly decreased levels of Arg8 in cells lacking Qcr8 and Cyt1, while absence of the late assembling Rip1 did not change Arg8 levels. Intriguingly, deletion of *MRX4* in these strains restored Arg8 synthesis to wild-type levels, demonstrating that absence of Mrx4 disrupts the *COB* translational feedback loop.

### Mrx4 interacts with the Cyt*b*-binding site of Cbp3

To investigate whether Cbp3 directly interacts with Mrx4, we used chemical crosslinking in intact mitochondria containing ALFA-tagged protein followed by purification and analysis with mass spectrometry (ALFA-XPL) (**Fig 4a)**. This confirmed the established interactions of Cbp3 to its partner protein Cbp6 and to the PTE proteins uL29m (Mrpl4) and uL24m (Mrpl40) ^19^, but also revealed a direct binding of Cbp3 to Mrx4 (**Fig 4b**).

**Figure 4:**
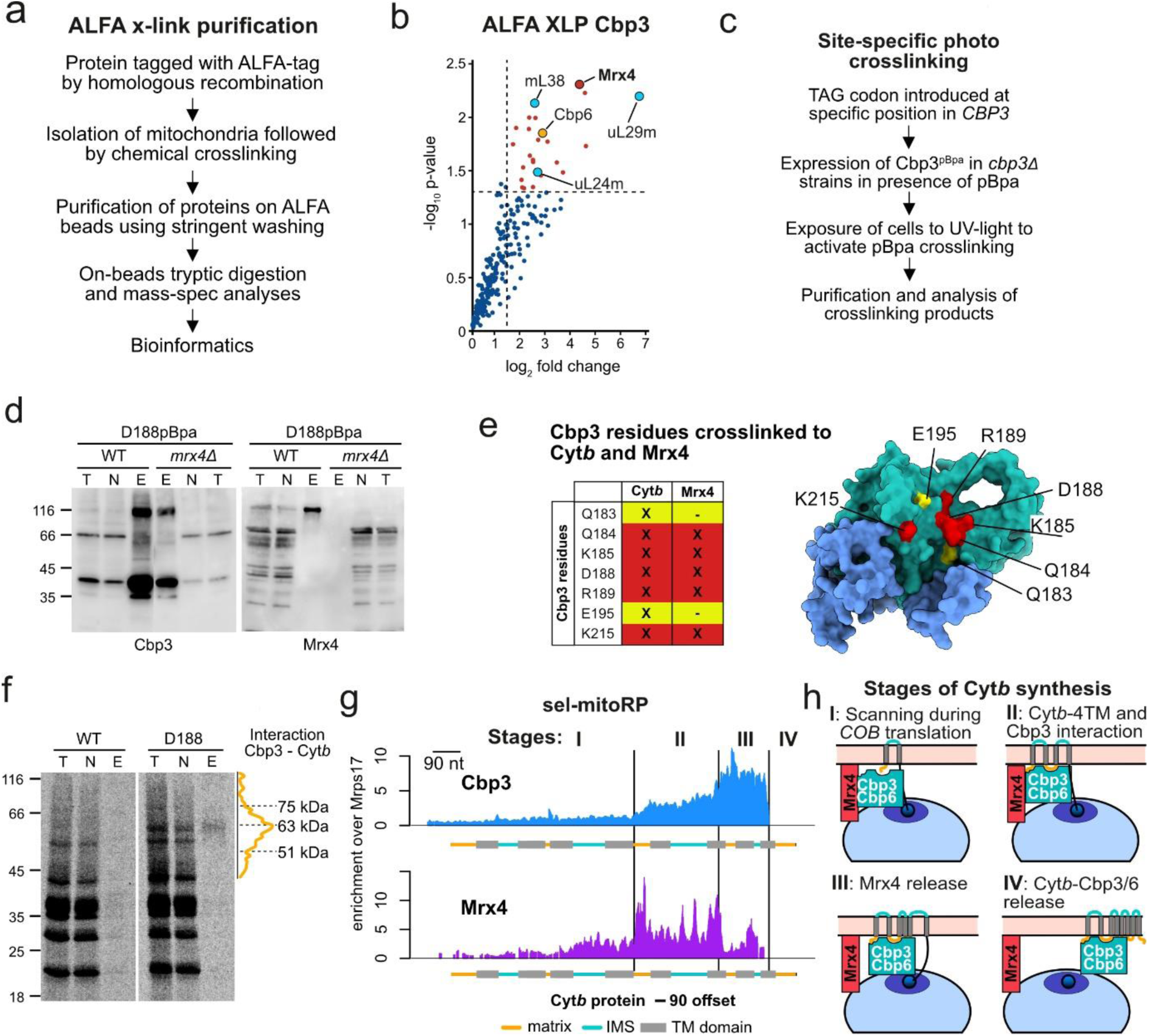
Mrx4 coordinates activating and repressive signals at the PTE. **(A)** Workflow for the stringent purification of proteins with a genomically integrated ALFA-tag following chemical crosslinking and with stringent washing. Samples were subsequently analyzed via on-bead tryptic digestion and mass spectrometry. **(B)** ALFA crosslinking purification (ALFA XLP) of Cbp3-ALFA followed by analysis with mass spectrometry. Data presented as log2 fold change compared to a Cbp3-ALFA control without crosslinking and with a threshold set to 1.5 (*n* = 3 for both conditions). **(C)** Scheme of the experimental approach for site-specific photo-crosslinking of Cbp3 by incorporation of the photoreactive amino acid p-benzoyl-L-phenylalanine (pBpa) at specific positions. Crosslinking was followed by Ni-NTA purification and analysis via Western blotting. **(D)** Site-specific photo-crosslinking and purification of Cbp3 with pBpa incorporated at residue D188 in the presence or absence of *MRX4*. Decoration with antibodies against Cbp3 and Mrx4 show that Cbp3 can be crosslinked to Mrx4 at this specific position. T = total, N = non-bound, E = elution. **(E)** Left: Table showing Cbp3 residues that previously have been photo-crosslinked to Cyt*b* (yellow) ^36^and now also to Mrx4 (red). Right: Mapping of these residues on the *AlphaFold2*-predicted structure of Cbp3-Cbp6 (Cbp3 in green and Cbp6 blue) points to a shared binding site of Cyt*b* and Mrx4 on the conserved chaperone domain of Cbp3. **(F)** Densitometric quantification of the smeary nascent Cyt*b* crosslinked to Cbp3 after *in-organello* labelling with 35S-methionine followed by site-specific photo-crosslinking and purification of residues Q184, D188 (visualized) and K215. T: total, N: non-bound, E: elution. **(G)** Selective mitoribosome profiling (sel-mitoRP) of Cbp3- and Mrx4-bound mitoribosomes demonstrate enrichment during the later stages of *COB* translation. Data presented as enrichment over mitoribosomes purified via Mrps17-3xFLAG. Cyt*b* is depicted with a 90 nt offset to represent emergence through the polypeptide tunnel. **(H)** Illustration of the four suggested stages of Cyt*b* synthesis with regards to interactions of Cbp3-Cbp6 (green) with Mrx4 (red) and the Cyt*b* nascent chain.

To further characterize the interaction between Cbp3 and Mrx4, we utilized site-specific photocrosslinking using the photoreactive amino acid p-benzoyl-L-phenylalanine (pBpa) incorporated at specific sites in Cbp3 ^36^. After exposure to UV-light, pBpa will crosslink to neighbouring residues yielding a covalent protein-protein crosslinking product that can be analyzed through purification and Western blotting (**Fig 4c)**. Using this approach, we have previously mapped the substrate binding site of Cbp3 ^36^, which binds to Cyt*b* and is formed by a V-shaped helix-turn-helix motif. Strikingly, the Cbp3 substrate binding site also formed specific crosslinks with Mrx4 at five residues (D188pBpa, Q184pBpa, K185pBpa, R189pBpa and K215pBpa) (**Fig 4d and e**). Therefore, Mrx4 binds directly to the substrate-binding site of Cbp3-Cbp6, an interaction that should be dissolved by binding of nascent Cyt*b* to Cbp3.

To reveal an interaction between nascent Cyt*b* and Cbp3, we used site-specific photocrosslinking to the substrate binding site in combination with labelling of mitochondrial translation products with ^35^S-Met. Purification of Cbp3 after crosslinking resulted in a specific signal of full-length Cyt*b* and a lower-molecular weight smear reflecting nascent Cyt*b* interacting with Cbp3 (**Fig 4f** and Ndi *et al.* 2019 ^36^). To determine the exact timing of the interactions between Cbp3 and Mrx4 with the mitoribosome during Cyt*b* synthesis, we performed sel-mitoRP on cells expressing Flag-tagged Cbp3 or Mrx4, respectively (**Fig 4g**). While only low intensity signals were recorded during early stages of *COB* translation, increased interactions were observed at a stage where the first four TMs of Cyt*b* have emerged from the ribosome. This signal was even further intensified at late stages of Cyt*b* synthesis. Mrx4 sel-mitoRP revealed a similar trend, but the signal of Mrx4 strongly declined upon emergence of the sixth TM of Cyt*b*. Taken together, these data reveal four different stages during Cyt*b* synthesis (**Fig 4h**): A first step (I), where Cbp3-Cbp6 bound to Mrx4 scans the PTE for emergence of the nascent Cyt*b*; a second step (II), where Cbp3-Cbp6 establishes an interaction with the portion of Cyt*b* encompassing the first four TMs. In the next step (III), the interaction between Mrx4 and Cbp3 is dissolved around emergence of the sixth TM segment. Finally (IV), the complex containing fully synthesized Cyt*b* and Cbp3/6 leaves the ribosome upon completion of translation.

### Mrx4 orchestrates the *COB* translational feedback loop at the PTE

Having identified the timing of Cytb-Cbp3/6 interaction, we next aimed to unravel how absence of Mrx4 can disrupt the regulation of *COB* translation. ALFA-XLP demonstrated that Mrx4 binds to Cbp3-Cbp6, the assembly factor Bca1 and, intriguingly, also to the *COB* TA Cbs2 (**Fig 5a**). Because binding of Cbs2 to the PTE coincides with repression of Cyt*b* synthesis ^17^, while binding of Cbp3-Cbp6 to the PTE activates translation ^20^, we hypothesized that the interaction of these factors with Mrx4 might be central to orchestrating the feedback loop by reciprocal interactions.

**Figure 5:**
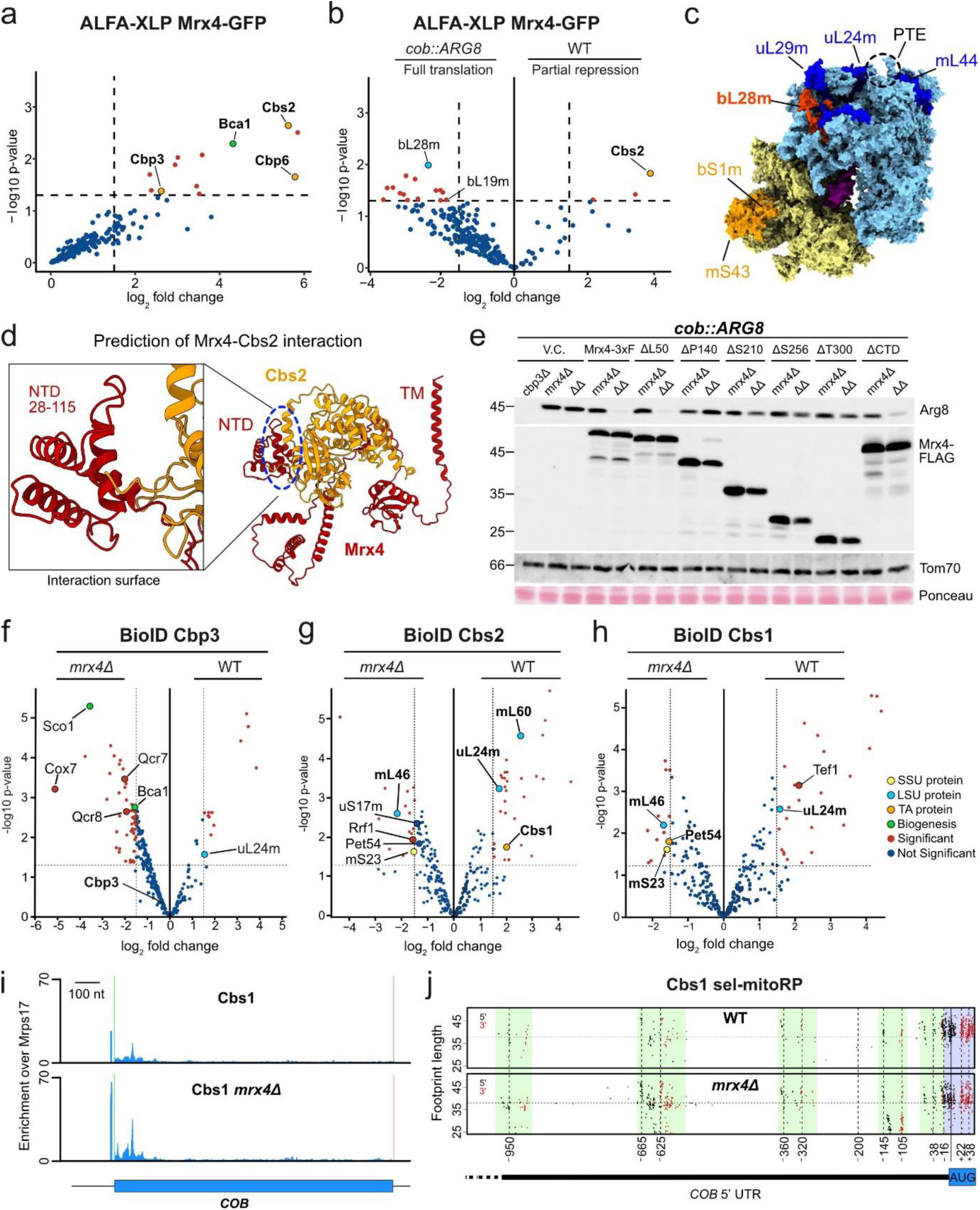
*COB* TAs shuttle the mRNA between the LSU and SSU. **(A)** Purification of Mrx4-GFP-ALFA from isolated mitochondria treated with the chemical crosslinker MBS (ALFA XLP). Data presented as log2 fold change compared to a control treated with DMSO and with a threshold set to 1.5 (*n* = 3 for both conditions). **(B)** ALFA XLP of Mrx4-GFP-ALFA in wild-type compared to the *cob::ARG8* genetic background, reflecting partial repression and full activation of *COB* translation, respectively. **(C)** Illustration of the location of mitoribosomal proteins on the structure. **(D)** *Alphafold2* prediction of Mrx4 (red) in complex with Cbs2 (orange). The main interaction is predicted to occur on a NTD of Mrx4 spanning residues 52-115. TM: transmembrane helix. NTD: N-terminal domain. **(E)** Steady-state levels of Arg8 in *cob::ARG8 mrx4Δ* and *cbp3Δmrx4Δ* strains expressing truncated versions of Mrx4-3xFLAG to test repressor function of Mrx4. The N-terminal was replaced by an Oxa1 MTS at indicated residues and for the CTD mutant a 3xFLAG tag was inserted before the TM helix. V.C. = vector control. Proximity labelling of **(F)** Cbp3-BirA*, **(G)** Cbs2-BirA* and **(H)** Cbs1-BirA* in the presence (WT) or absence of Mrx4 (*mrx4Δ*) with data presented as significant with a log2 fold change over 1.5 and a log10 p-value under 0.5. **(I)** Selective mitoribosome profiling (sel-mitoRP) of Cbs1-bound mitoribosomes shows enrichment during initiation of *COB* translation, which is markedly increased (2.5x) in cells lacking *MRX4*. Both sets of data are presented as enrichment over mitoribosomes purified via Mrps17-3xFLAG, in wild-type or *mrx4Δ* strain backgrounds, respectively. **(J)** V-plot of protected footprints at specified positions in the 5’-UTR of *COB* mRNA after sel-mitoRP of Cbs1-bound mitoribosomes in wild-type and *mrx4Δ* strains. The five main regions of footprints (marked in green) all show more reads in the strain lacking Mrx4, indicating a higher occupancy of *COB* TAs. The region with footprints protected by the mitoribosome is highlighted in blue.

To test this, we compared interaction patterns of Mrx4 in cells with fully active translation of *COB* mRNA (using a strain with the *cob:ARG8* mtDNA lacking Cyt*b*, hence without sequestration of Cbp3-Cbp6 in assembly intermediates) with the wild type scenario, where *COB* translation is substantially repressed due to feedback inhibition ^21^. As predicted, full translation in the strain carrying *cob:ARG8* impaired the contact between Mrx4 and Cbs2, which was clearly enriched in wild-type mitochondria (**Fig 5b**). Moreover, Mrx4 contacted the LSU protein bL28m (Mrpl28), which localizes on the ribosome between the PTE and MCE, during full translation, suggesting that Mrx4 switches localization (and presumably changes conformation) in a *COB*-translation dependent manner. While under wild type conditions, Bio-ID revealed that Mrx4 is primarily bound to the PTE in vicinity to mL44 and Cbs2 (**Fig 2c**), the contact to Cbs2 is decreased and new contacts with bL28m are established during full *COB* translation (**Fig 5c**).

Structure prediction using AlphaFold2 suggested that Mrx4 interacts with Cbs2 with a small globular domain spanning residue 30-115 (**Fig 5d**), a contact presumably representing the repressed state of *COB* translation. To test whether repression indeed depends on this predicted interaction, we expressed truncated variants of Mrx4 in cells harboring the *cob::ARG8* mtDNA and lacking both Cbp3 and Mrx4 (**Fig 3b**). In this strain, the *cob::ARG8* mRNA was fully translated **(Fig 3c)**. Expression of full length Mrx4 or a mutant lacking the first 50 amino acids of the mature N-terminus restored repression, while expression of constructs lacking 140 or more residues of the N-terminus could not repress Arg8 synthesis (**Fig 5e**). These data indicate that Mrx4 contacts Cbs2 with a domain located between residues 50 and 140 to mediate translational repression of the *COB* mRNA.

Having identified Mrx4 as the key component of this translational feedback loop, we next sought to understand the localization dynamics of the *COB* TAs and assembly factor with and without Mrx4. Using BioID, we found that Cbp3 binding to the PTE (as evidenced by its proximity to uL24m) was enhanced in the presence of Mrx4, while Cbp3 increased proximity to respiratory chain assembly factors in the absence of Mrx4 (**Fig 5f**). Comparing the Cbs1 and Cbs2 proximity interactomes revealed differences in wild type and *mrx4*Δ cells, reflecting the switch from partial repression to full translation of *COB* mRNA (**Fig 5g**). In wild-type cells, these *COB* TAs were located close to the PTE (as evidenced by uL24m), revealing its interactions during the repressed state of *COB* mRNA translation. However, in the absence of Mrx4, they changed sub-ribosomal localization and were found at the SSU and the MCE. **(Fig 5g, h**). Consequently, sel-mitoRP of Cbs1 in the absence of Mrx4 revealed substantially increased reads in the 5′-end of the *COB* mRNA (**Fig 5i**), concomitant with increased footprints of protected fragments in the 5′-UTR (**Fig 5j**), indicating increased translation initiation upon disruption of the translational feedback loop.

Together, these data reveal that Mrx4 orchestrates a spatial reorganization of both *COB* translational activators and assembly factors, maintaining TAs in proximity to the PTE when translation needs to be restrained, while also ensuring proper PTE positioning of assembly factors for efficient complex III assembly. In the absence of Mrx4, these factors undergo distinct relocalizations - TAs shift to sites of translation initiation leading to enhanced Cyt*b* synthesis, while assembly factors show increased associations with other respiratory chain assembly components, likely reflecting their redirected activities when the feedback loop is disrupted. These findings establish Mrx4 as the molecular switch that controls this translational feedback loop through its ability to dynamically regulate the spatial distribution of both translational activators and assembly factors at the mitoribosome.

## Discussion

Mitochondrial translation is highly specialized to produce a handful of mostly very hydrophobic proteins. The small number of protein-coding genes in mtDNA facilitated the establishment of mechanisms that allow for a tailored biogenesis of its translation products, including highly specialized mechanisms for translational regulation. The coordination of mitochondrial and nuclear gene expression is a key challenge during the biogenesis of the OXPHOS system, which is solved in the case of yeast Cyt*b* through a translational feedback loop targeting translation initiation of the *COB* mRNA. Here, we have identified how Mrx4, a ligand of the mitoribosomal PTE, acts as a molecular switch in this process. Mrx4 provides a binding site for *COB* mRNA complexed with two translational activators, which is necessary to repress its translation. The presence of the assembly factor Cbp3-Cbp6 at the PTE and interaction with Mrx4 activates the switch, triggering the release of the *COB* TA complex, which can then interact with the MCE for translation initiation. Through this mechanism, the presence of Cbp3-Cbp6 at the PTE can be sensed and integrated into a signal for enhanced Cyt*b* synthesis.

Translational repression in the cytosol works by various strategies including sequestering mRNAs into higher-order structures to exclude these mRNAs from translation initiation ^37,38^, by inhibiting key initiation factors by direct interactions or through posttranslational modifications ^39,40^, or through mechanisms of RNA interference ^41^. Organellar protein synthesis depends on and reacts to nuclear gene expression ^1,2^, but also maintains mechanisms for translational regulation ^7,13,42^. For this, endosymbiotic organelles like mitochondria and chloroplasts contain specific RNA binding proteins ^43,44^, which often contain pentatricopeptide repeat (PPR) motifs to recognize specific mRNAs. Many of these RNA-binding proteins thereby interact with one specific mRNA, often in 5′-UTRs, and are necessary to activate translation of this mRNA. Work in yeast has shown that these translational activators bind to the MCE ^18,28^ to help align the start codon into the ribosomal P-site, thereby replacing the missing Shine-Dalgarno sequence as molecular guides ^28^. Protected fragments of the 5′-UTR of *COB* mRNA revealed in this work suggest that the TAs bind this mRNA in a folded state that spans large portions of the mRNA, similar to the case of the *ATP9*-Aep1-Aep2-Atp25C complex ^28^. Specifically, the portion around 100 nucleotides relative to the start codon is likely important for translation initiation, as this part should determine start codon alignment. Previous genetic mapping ^29^ revealed that this region of the 5’-UTR is bound by Cbs2, which in turn binds to the MCE. Hence, it is conceivable that Cbs2 plays a dedicated role for aligning the *COB* mRNA into the mRNA channel, while Cbs1 and Cbp1 could impose or stabilize a specific fold of the 5′-UTR, which would require a certain flexibility to allow for the observed translocations of the TA complex from the MCE to the PTE and *vice versa*.

The PTE is the site of early protein biogenesis. Upon emergence from the tunnel exit, early folding events and protein maturation occur that are often mediated by specific proteins including chaperones, membrane insertases and processing enzymes ^27,45^. These early biogenesis factors interact with ribosomal proteins decorating the surface surrounding the PTE. Here, we have identified Mrx4 as a novel ligand of the mitoribosomal PTE that does not play a role in early protein biogenesis but is necessary for translational regulation of a single mRNA. To this end, Mrx4 interacts reciprocally with either the *COB*-TA complex or the Cyt*b*-specific chaperone Cbp3-Cbp6. Interestingly, Mrx4’s crosslinking partners were altered during fully active translation, which would be in line with a conformational change upon loss of interaction with *COB*-TA and binding to Cbp3-Cbp6. Strikingly, both Mrx4 and Cyt*b* engage with the substrate binding site of Cbp3. Hence, the interaction between Cbp3-Cbp6 and Cyt*b* apparently competes off Cbp3-Cbp6 from Mrx4, revealing the time point when the translational switch is activated. Accordingly, a plausible model for *COB* translational control (**Fig 6**) is that Mrx4 binds the *COB* TAs. Once Cbp3-Cbp6 detaches from Cyt*b*, the interaction of this assembly factor with Mrx4 releases the *COB* TAs. The TAs, bound to *COB* mRNA, are then free to localize to the MCE, enabling translation initiation. Early during elongation, the TAs leave the MCE and the mitoribosome. Meanwhile, Mrx4-bound Cbp3-Cbp6 scans the nascent Cyt*b* at the PTE. Upon emergence of the loop after the fourth TM segment from the ribosome, Cbp3-Cbp6 binds the nascent Cyt*b*. Subsequently, Cbp3-Cbp6 is released from Mrx4 (when the sixth TM segment emerged from the tunnel), upon which Mrx4 is available to “reload” through interacting with the *COB*-TA complex and inhibit further translation initiation on *COB*.

**Figure 6:**
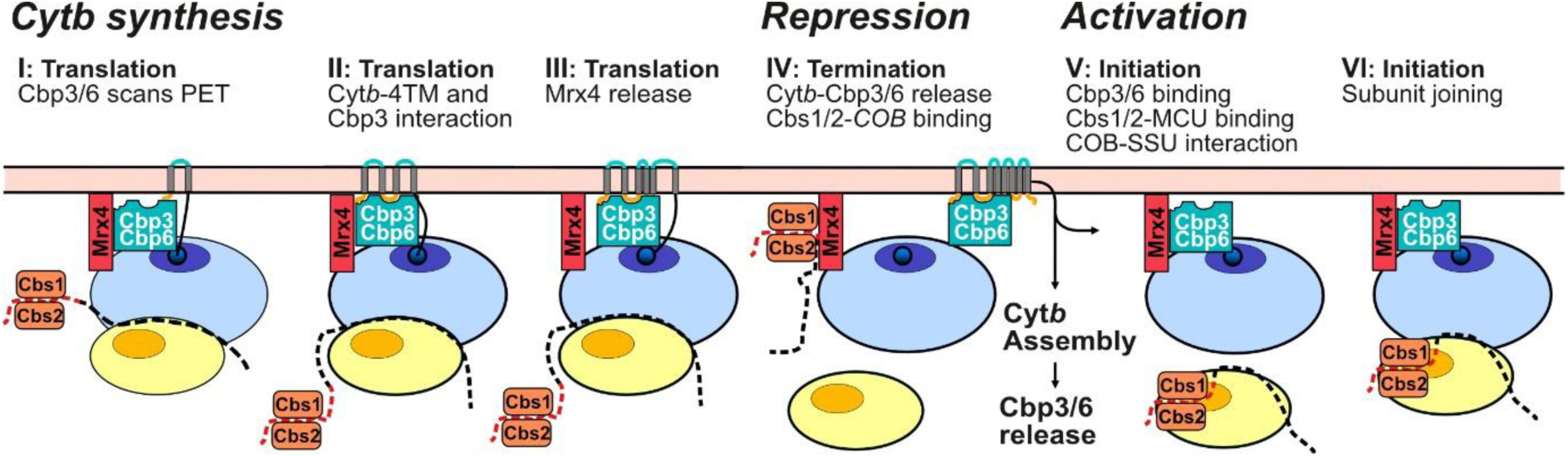
Orchestration of Cyt*b* biogenesis and translation control at the mitoribosomal PET. A model of the translational feedback loop and dynamics of protein biogenesis of the mitochondrial encoded Cyt*b*. Cyt*b* biognesis relies on distinct steps of protein interactions to during *COB* translation to ensure that newly synthesized Cyt*b* can be bound to its dedicated chaperone Cbp3-Cbp6. Once Cyt*b* is fully synthesized, it leaves the mitoribosome in a complex with Cbp3-Cbp6. Through this, the PTE ligand Mrx4 is available to bind the *COB* TA Cbs2, thereby sequestering the *COB* TA complex at the PTE. Release of Cbp3-Cbp6 from Cyt*b* during assembly allows to activate a new round of *COB* translation. To this end, Cbp3-Cbp6 competes off The *COB* TA complex from Mrx4. The thereby released *COB* TA complex can now interact with the MCE at the SSU for translation initiation.

Finally, a complex containing full-length Cyt*b* and Cbp3-Cbp6 detaches from the mitoribosome to mediate Cyt*b* maturation and assembly. Once Cyt*b* is fully hemylated and interacts with the first nuclear encoded subunits ^21^, Cbp3-Cbp6 is released from Cyt*b* to activate a new round of *COB* translation.

In addition to *COB* mRNA, *VAR1*, *COX1* and *ATP6*, coding for key subunits of the small mitoribosomal subunit, cytochrome *c* oxidase or ATP-synthase, respectively, are subject to translational regulation in yeast mitochondria ^31,34,35,46^. In these cases, sequestration of the translation products in assembly intermediates represses new rounds of translation, but how this is relayed to translation is yet unknown. Cox14 is a cytochrome *c* oxidase assembly factor that stabilizes the Cox1-containing assembly intermediate important for translational repression ^35,47,48^. In its absence, this assembly intermediate cannot form efficiently and hence sequestration of the regulating TA, Mss51, in the intermediate is less efficient, leading to a restoration of *COX1* translation. Interestingly, the protein Smt1 (Mrx5) was recently identified as a repressor of Atp6 and Atp8 synthesis ^49^ that binds the bicistronic *ATP8/ATP6* mRNA. How Smt1 mediates this repression is yet unknown. However, given that Smt1 interacts with mitoribosomes ^33^, it is possible that it operates through a similar mechanism as Mrx4. It will be an exciting task for future research to identify these molecular mechanisms that will likely show compelling variations of translational regulation in mitochondria.

## Material and methods

### Yeast strains used in the study

All strains in this study were isogenic to *Saccharomyces cerevisiae* strains W303 *MAT*a {*leu2-3,112 can1-100 trp1-1 ura3-1 ade2-1 his3-11,15*} or S288c ^1^ and are listed in Table XXX. Strains with modified mitochondrial genomes were created in previous studies ^20,34^. Chromosomal open reading frames (ORFs) were modified via homologous recombination according to ^50^. ORFs were disrupted by insertion of *KanMX4*, *URA3*, *LEU2* and Hygromycin (hph) selection cassettes. FLAG, ALFA and BirA* C-terminal epitope tags were added by replacing the stop codon of endogenous ORFs with the respective tag sequence followed by *HIS3* or *TRP1* selection cassettes. After transformation, genomic modifications were confirmed by growth on selection media and control PCR. For quantitative experimental approaches, 3 to 5 positively tested clones were applied to rule out congenic variations.

### Media and culturing conditions

Strains were grown at 30°C and 170 rpm in either full media containing 1% yeast extract, 2% peptone and 2% glucose (YPD), galactose (YPGal) or glycerol (YPG) as indicated, or synthetic complete (SC) media, consisting of 0.17% yeast nitrogen base, 0.5% (NH_4_)_2_SO_4_ and 30 mg/l of all amino acids (except 80 mg/l histidine and 200 mg/l leucine) and 30 mg/l adenine with 2% galactose.

### Mitochondrial isolation

Yeast cells were grown overnight in YPG or YPGal to exponential phase (OD_600_ = 1-3) before harvesting through centrifugation at 3,000x*g* for 5 min. Cells were washed with distilled water, resuspended in MP1 buffer (2 ml/g wet weight of 0.1 M Tris-base, 10 mM dithiothreitol (DTT) according to 2 ml/g wet weight), incubated for 10 min at 30°C before being washed with 1.2 M sorbitol, resuspended in MP2 buffer (6.7 ml/g wet weight if 20 mM KPi pH 7.4, 0.6 M sorbitol and 3 mg/g wet weight of zymolyase (Seikagaku Biobusiness, Tokyo, Japan) and incubated at 30°C for 1 h to digest the yeast cell wall. Spheroplasts were harvested at 3000xg for 5 min at 4°C, resuspended in MP3 buffer (13.4 ml/g wet weight of 0.6 M sorbitol, 10 mM Tris pH 7.4, 1 mM EDTA and 1 mM PMSF) and homogenized in 2 x 10 strokes using a tight-fitting homogenizer (Sartorius Stedim Biotech S.A., France). The homogenate was centrifuged two times at 3000xg for 5 min at 4°C before mitochondria were harvested by centrifugation at 15.000xg for 15 min at 4°C. The mitochondrial pellet was resuspended in SH buffer (0.6 M sorbitol and 20 mM HEPES pH 7.4) to a final concentration of 10 mg/ml and frozen in liquid nitrogen before storage at -80°C.

### Carbonate extraction and protease protection assay

For the carbonate extraction assay, 100 µg of isolated mitochondria was resuspended in 0.1 M Na_2_CO_3_ (treatment) or 0.1 M NaCl (control), incubated for 30 min on ice, and centrifuged for 100,000 x *g* for 30 min at 4°C. Membrane and soluble fractions were TCA precipitated and analyzed by SDS-PAGE and Western blotting. For the protease protection assay, 100 µg of isolated mitochondria was incubated in SH buffer (whole mitochondria), 20 mM HEPES/KOH pH 7.4 (mitoplasts) or lysis buffer (SH buffer with 0.2% Triton X-100) for 30 min on ice. Proteinase K was added to a final concentration of 10 µg/ml and incubated for 20 min on ice. PMSF was added to a final concentration of 1 mM before intact mitochondria and mitoplasts were centrifuged at 25,000 x *g* for 10 min at 4°C. Supernatants and pellets were TCA precipitated and analyzed by SDS-PAGE and Western blotting.

### In-vivo and in-organello radiolabelling

*In-vivo* labelling of mitochondrially encoded translation products was performed according to the same procedure as previously described ^51^. Shortly, cells in logarithmic growth phase (OD600 1-2) were washed in SGal media without amino acids, after which they were resuspended in SGal with all amino acids lacking methionine. After an incubation time of 5 min at 30°C, ^[35]^S-labeled methionine was added and aliquots were taken after 5, 10 and 15 min. Unlabeled methionine was added [10 mM] and the temperature increased to 37°C. Aliquots were taken after 30, 60 and 90 min and proteins were directly extracted with TCA precipitation and analyzed with SDS-PAGE and autoradiography.

### Analysis of mitoribosome interaction via linear sucrose gradients

Isolated mitochondria were suspended in lysis buffer (10 mM Tris-HCl pH 7.4, 10 mM KOAc, 0.5 mM Mg(OA)_2_, 10 mM EDTA, 5 mM b-mercaptoethanol (BME), 1% n-dodecyl-b-D-maltoside (DDM), 1 mM phenylmethylsulfonyl fluoride (PMSF), 1x complete protease inhibitor (Roche), 0.1 mM spermidine and 5% glycerol) and lysed for 10 min at 4°C. Lysate was diluted with one volume of lysis buffer without DDM, centrifuged at 16,000 x *g* for 10 min at 4°C, and loaded on a linear sucrose gradient (0.3 – 1 M sucrose in lysis buffer) which was centrifuged at 60,000 rpm in a SW60 rotor for 1 h at 4°C. Collected fractions were TCA precipitated and pelleted proteins were washed twice with acetone and resuspended in sample buffer followed by analysis through SDS-PAGE and Western blotting.

### Ribosome sedimentation via sucrose cushion

Mitochondria were lysed in lysis buffers of varying ionic strength (10 mM Tris-HCl pH 7.4, 10/50/100/300 mM KOAc, 2.5/5/20/50 mM Mg(OAc)_2_, 0.8 mM EDTA, 1 % DDM, 5% glycerol, 5 mM BME, 1 mM PMSF, 1 mM spermidine, 1x Complete protease inhibitor (Roche) for 10 min on ice. Lysate was diluted in one volume of respective lysis buffer without DDM and clarified by centrifugation at 16,000 x *g* for 10 min at 4°C. One half of the cleared lysate was TCA precipitated as a Total sample (T) while the other half was loaded on a sucrose cushion (1.2 M sucrose in lysis buffer) and centrifuged at 190,000 x *g* for 105 min at 4°C. Supernatant (S) and pellet (P) were both TCA precipitated and all samples were separated by SDS-PAGE and analyzed by Western blotting.

### Site-specific photo-crosslinking

Site-specific photo-crosslinking of the conserved chaperone domain of Cbp3 was performed as previously described ^36^.

### Chemical crosslinking and affinity purification

Mitochondria (5 or 10 mg) from yeast strains expressing protein of interest with a C-terminal ALFA or GFP-ALFA epitope tag were suspended in SH buffer (0.6 M sorbitol and 20 mM HEPES pH 7.4) and incubated for 5 min at 30°C. Membrane permeable chemical crosslinkers m-maleimidobenzoyl-N-hydroxysuccinimide (MBS) or bismaleimidoethane (BMOE) dissolved in dimethyl sulfoxide (DMSO) were added to a final concentration of 200 µM and samples were incubated for 30 min at 30°C, 600 rpm. DMSO only was added as non-crosslinked control. Crosslinking was stopped by addition of 100 mM Tris-HCl pH 8 and 100 mM BME followed by incubation for 10 min at 30°C. Mitochondria were pelleted through centrifugation at 25,000 x *g* for 10 min at 4°C and lysed in lysis buffer (20 mM Tris-HCl pH 7.4, 150 mM KCl, 1% DDM, 1 mM PMSF, 1x Complete protease inhibitor (Roche) and 8 M Urea) for 30 min, tumbling at room temperature. One volume of dilution buffer (20 mM Tris-HCl pH 7.4, 150 mM KCl, 0.01% DDM) was added and sample was clarified through centrifugation at 16,000 x *g* for 10 min at 4°C. Clarified lysate was added to 40 µl slurry of ALFA selector ST beads (NanoTag Biotechnologies, Göttingen, Germany), incubated for 2 h at room temperature, washed 3 times with wash buffer A (20 mM Tris-HCl pH 7.4, 0.1 % SDS), 3 times with wash buffer B (20 mM Tris-HCl pH 7.4, 6 M Urea) and 3 times with ABC buffer (50 mM ammonium bicarbonate pH 8.0) before resuspension in ABC buffer containing 0.5 µg sequencing grade trypsin (Promega). After on-bead tryptic digestion overnight at 37°C, the supernatant was harvested before addition of formic acid to a final concentration of 1%. Peptides were lyophilized for further analysis by mass spectrometry.

### Proximity labelling (Bio-ID)

Proximity labelling via Bio-ID was performed as previously described ^18^. In short, yeast strains with protein of interest containing a C-terminal BirA* tag was grown overnight in YPG or YPGal supplemented with 50 µM biotin. The next day, mitochondria were isolated (see section “Isolation of mitochondria”) in three biological replicates per strain. Mitochondria (3 mg) were lysed in 1% SDS at 50°C for 5 min, diluted in RIPA buffer (50 mM Tris-HCl pH 7.5, 150 mM NaCl, 1% NP-40, 1 mM EDTA, 1 mM EGTA, 0.1% SDS, 0.5% sodium deoxycholate, 1x Complete protease inhibitor (Roche) and 1 ul of benzonase (2U, Sigma Aldrich) and incubated for 30 min on ice. Lysate was clarified by centrifugation for 10 min at 16,000 x *g* at 4°C and added to pre-equilibrated streptavidin magnetic beads (Thermo Fischer) for 3 hours at 4°C. Beads were washed 2 x with RIPA buffer, 2 x with TAP lysis buffer (50 mM HEPES pH 8, 100 mM KCl, 10 % glycerol, 2 mM EDTA and 0.1 % NP-40), 3 x with ABC buffer (50 mM ammonium bicarbonate pH 8). Biotinylated proteins were on-bead-digested with 1 µg sequencing grade trypsin (Promega) overnight at 37°C and peptides were lyophilized and analyzed by mass spectrometry.

### Mass spectrometry

For analysis with LC-MS/MS, desalted peptides were injected in an Ultimate 3000 RSLCnano system (Thermo), separated in a 15-cm analytical column (75 mm ID home-packed with ReproSil-Pur C18-AQ 2.4 mm from Dr. Maisch) with a 50-min gradient from 5 to 60% acetonitrile in 0.1% formic acid. The effluent from the HPLC was directly electrosprayed into a Qexactive HF (Thermo) operated in data dependent mode to automatically switch between full scan MS and MS/MS acquisition. Survey full scan MS spectra (from m/z 375–1600) were acquired with resolution R = 60,000 at m/z 400 (AGC target of 3x106). The 10 most intense peptide ions with charge states between 2 and 5 were sequentially isolated to a target value of 1x105, and fragmented at 27% normalized collision energy. Typical mass spectrometric conditions were: spray voltage, 1.5 kV; no sheath and auxiliary gas flow; heated capillary temperature, 250_C; ion selection threshold, 33.000 counts. MaxQuant 1.5.2.8 was used to identify proteins and quantify by iBAQ with the following parameters: Database, UP000002311_559292_Scerevisiae_20171017; MS tol, 10ppm; MS/MS tol, 0.5 Da; Peptide FDR, 0.1; Protein FDR, 0.01 Min. peptide Length, 5; Variable modifications, Oxidation (M); Fixed modifications, Carbamidomethyl (C); Peptides for protein quantitation, razor and unique; Min. peptides, 1; Min. ratio count, 2.

### Selective mitoribosome profiling

Selective mitoribosome profiling was carried out as described previously (Bridgers and Carlstrom et al). Briefly, yeast were grown in YPGal (1% yeast extract, 2% peptone, 2% galactose), pH 5.0, until an OD600 of approximately 1. 150 mL of culture was snap chilled over ice prior to formaldehyde crosslinking. The crosslinking was quenched with glycine and washed twice in 40 mL of ice cold crosslinking wash buffer (50 mM HEPES pH 7.5, 50 mM NH4Cl, 10 mM MgCl2). The pellet was then resuspended in 4 mL of crosslinking lysis buffer (50 mM HEPES pH 7.5, 50 mM NH4Cl, 10 mM MgCl2, 1.5X Complete EDTA-free protease inhibitor cocktail, 0.5% lauryl maltoside) and dripped into a 50 mL conical filled with liquid nitrogen to form frozen droplets.

The frozen cells with lysis buffer were then mechanically lysed using liquid nitrogen-chilled 50 mL canisters for six cycles of 3 min at 15 Hz using a Retsch MM301 mixer mill. The thawed cell lysate was diluted with 2 mL of fresh crosslinking lysis buffer and treated with 300 Units of RNase I (Epicentre) for 30 minutes in a room temp water bath, swirling halfway to mix. 600 units of SUPERase IN RNase inhibitor was added to the lysate to stop RNase digestion. The clarified lysate was layered on top of 8 mL of sucrose cushion buffer (50 mM HEPES pH 7.5, 50 mM NH4Cl, 10 mM MgCl2, 1.5X Complete EDTA-free protease inhibitor cocktail, 24% sucrose) and then centrifuged for 4.5 hours at 40,000 rpm at 4 deg C.

The ribosome pellet was resuspended in chilled mitoribosome wash buffer (50 mM HEPES pH 7.5, 50 mM NH4Cl, 10 mM MgCl2, 0.1% Triton-X-100) overnight at 4 deg C with shaking. The resuspended ribosome pellet was then added to 50 uL of packed Anti-FLAG M2 Affinity Gel and incubated with end over end rotation for 3 hrs at 4 deg C. The resin was then washed 3 times with 1 mL of cold mitoribosome wash buffer. Following the last wash, the resin was resuspended with 1 mL of mitoribosome wash buffer using a cut pipette tip and transferred to a fresh 1.5 mL tube. The wash was removed and the resin was resuspended in 600 uL of mitoribosome wash buffer with 200 ug/mL 3x-FLAG peptide. The resin was then incubated for 1 hr at 4 deg C with rotation. To recover the elution, the resin slurry was passed over a Co-Star SpinX column for 1 min at 16,000 X G. The elution was then transferred to a fresh 1.5 mL tube. To reverse the formaldehyde crosslinking, 30 uL of 20% SDS, 32 uL of 100 mM DTT, and 14 uL of 0.5 M EDTA were added to the elution, which was then heated for 45 min at 70 deg C. The RNA footprints were extracted following reversal of the crosslinking using an equal volume of acid-phenol chloroform. Ribosome footprints were then isolated, and a cDNA library was prepared for Illumina short read sequencing as previously described ^52^.

### Selective ribosome profiling data analysis

Selective ribosome profiling data analysis was performed as described in (Bridgers and Carlstrom et al). Briefly, reads were trimmed, filtered for abundant non-coding RNA, and aligned to the *S. cerevisiae* genome assembly R64 (UCSC: sacCer3) for S288C strain background or assembly ASM216351v1 for W303 strain background. Ribosome A site positions were determined using an offset from the 3’ end of each read, depending on its length: 37:[-15], 38:[-16], 39:[-17], 40:[-17], 41:[-17]. Enrichment was calculated after combining replicates by dividing by the rpm values in a sliding 9-nt window in the corresponding total (Mrps17) dataset. The used scripts are available in Github (https://github.com/churchmanlab/Yeast_selective_mitoRP).

## Acknowledgements

We would like to thank all members of our research groups for stimulating discussions. This work was supported by the Swedish Research Council (2014-4116 and 2018-03694 to MO), the Knut and Alice Wallenberg foundation (2017.009 and 2019.0319 to MO) and the National Institutes of Health grant R01-GM123002 (L.S.C.). Proteomic analysis was performed at the Protein Analysis Units (ZfP) of the Ludwig Maximilians University of Munich, a registered research infrastructure of the Deutsche Forschungsgemeinschaft (DFG: 453504869).

## Contributions

A.C. generated strains and constructs, performed most protein analyses and prepared the first draft of the manuscript. J.B.B., M.T.C and A.C. generated the FLAG-tagged yeast strains for sel-mitoRP. J.B.B. performed the sel-mitoRP for Mrx4 and Cbp3. J.B.B. and M.T.C. analyzed the sel-mitoRP sequencing results. A.S. participated in Bio-ID data analyses, while I.F. and A.I. performed mass-spectrometry. M.O. and L.S.C. supervised the project, interpreted the data and wrote the paper. All authors contributed toward the final version of the paper.

