## Supplemental Figure 1 for "A molecular switch at the yeast mitoribosomal tunnel exit controls cytochrome *b* synthesis"

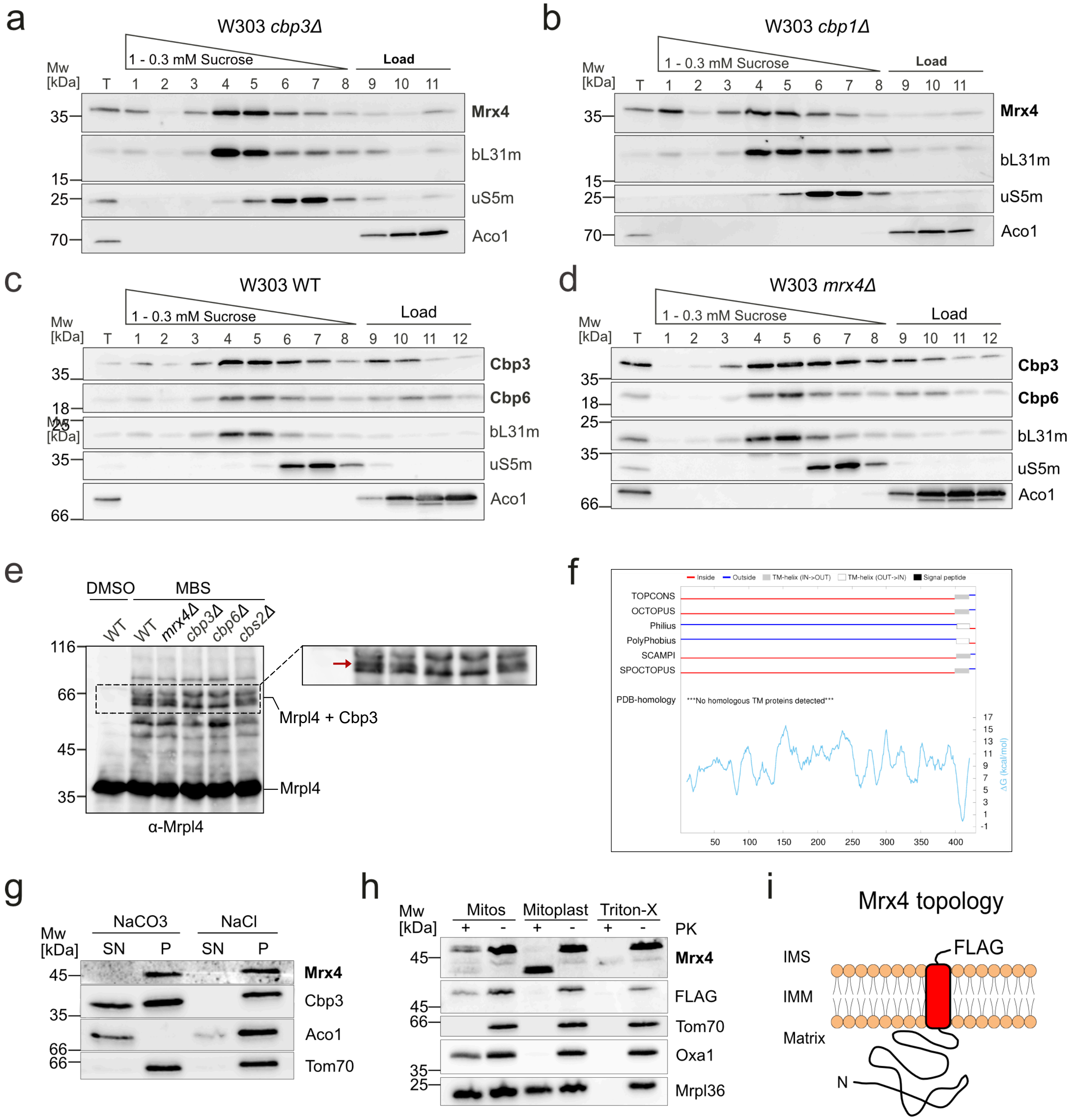


**Supplemental figure 1: (a)** Sucrose density gradients using mitochondria, where *CBP3* or *CBP1* **(b)** have been deleted, show no effect on Mrx4 co-migration with the mitoribosome. **(c)** Sucrose density gradients demonstrating that co-migration of Cbp3-Cbp6 with the mitoribosome does not change in the presence or absence of Mrx4. **(d)** *In situ* chemical crosslinking reveals binding of Cbp3 to the PTE in intact mitochondria in the absence of Mrx4. Mitochondria isolated from the indicated strains where exposed to the chemical crosslinker MBS, after quenching, proteins were separated on SDS-PAGE and analyzed with Western blotting against Mrpl4 (uL29m). A specific crosslinking product is formed that is absent in strains lacking Cbp3 or Cbp6, but not when Cbs2 or Mrx4 are deleted. **(f)**Topology prediction for Mrx4 (YPL168W) predicts Mrx4 to be an integral membrane protein with a single transmembrane helix close to the C-terminal end. **(g)** Mitochondria where exposed to alkaline treatment or high salt and separated into a membrane fraction (P) and a soluble fraction (SN). Mrx4 behaved as the predicted membrane proteins. **(h)** Mitochondria in isotonic media (Mitos), in hypotonic media (Mitoplast) or dissolved in detergent (Triton-X) were exposed to proteinase K (PK). Mrx4, carrying a C-terminal FLAG-tag, was protected from PK in intact mitochondria and cleaved into an N-terminal fragment in mitoplasts, establishing the topology depicted in **(i)**.
